# Systems Biology Analysis of Human Genomes Points to Key Pathways Conferring Spina Bifida Risk

**DOI:** 10.1101/2021.07.02.450913

**Authors:** Vanessa Aguiar-Pulido, Paul Wolujewicz, Alexander Martinez-Fundichely, Eran Elhaik, Gaurav Thareja, Alice AbdelAleem, Nader Chalhoub, Tawny Cuykendall, Jamel Al-Zamer, Yunping Lei, Haitham El-Bashir, James M. Musser, Abdulla Al-Kaabi, Gary M. Shaw, Ekta Khurana, Karsten Suhre, Christopher E. Mason, Olivier Elemento, Richard H. Finnell, M. Elizabeth Ross

## Abstract

Spina bifida (SB) is a debilitating birth defect caused by multiple gene and environment interactions. Though SB shows non-Mendelian inheritance, genetic factors contribute to an estimated 70% of cases. Nevertheless, identifying human mutations conferring SB risk is challenging due to its relative rarity, genetic heterogeneity, incomplete penetrance and environmental influences that hamper GWAS approaches to untargeted discovery. Thus, SB genetic studies may suffer from population substructure and/or selection bias introduced by typical candidate gene searches. We report a population based, ancestry-matched whole-genome sequence analysis of SB genetic predisposition using a systems biology strategy to interrogate 298 case-control subject genomes (149 pairs). Genes that were enriched in likely gene disrupting (LGD), rare protein-coding variants were subjected to machine learning analysis to identify genes in which LGD variants occur with a different frequency in cases vs. controls and so discriminate between these groups. Those genes with high discriminatory potential for SB significantly enriched pathways pertaining to carbon metabolism, inflammation, innate immunity, cytoskeletal regulation and essential transcriptional regulation, indicating their impact on the pathogenesis of human SB. Additionally, interrogation of conserved non-coding sequences identified robust variant enrichment in regulatory regions of several transcription factors critical to embryonic development. This genome-wide perspective offers an effective approach to interrogation of coding and non-coding sequence variant contributions to rare complex genetic disorders.

## Introduction

The neural tube defect (NTD) Spina Bifida (SB), among the debilitating but survivable malformations in live births, is due to failed embryonic neural tube closure. SB and the non-viable NTD, anencephaly, together have a global prevalence ranging from one in 3,000 to one in 100[73]. Decades of clinical and animal model investigations have indicated that SB comprises a complex genetic disorder, requiring at least one (and probably several) of many genetic alterations and/or gene-environment interactions for neurulation to fail[72, 84]. NTD-causing mutations have been reported in more than 250 mouse genes[29, 36], which has since grown to over 400 mutant genes currently listed in the Mouse Genome Informatics (MGI) database, further underscoring the complex genetic origins of the disorder. Genetic heritability of human SB, or the proportion of cases that are attributable to inherited genetic alteration, is estimated to be as much as 70%[44].

Maternal periconceptional supplementation with folic acid (vitamin B9) can reduce the occurrence of SB in offspring by as much as 70% in some populations[12, 21, 63]. Despite folate supplementation campaigns and fortification of the US food supply since 1998, SB prevalence rates have only dropped 30%, suggesting maximal benefit for folic acid has been achieved. Other agents such as vitamin B12, methionine or inositol show some promise for effective prevention[87]. However, the mechanisms through which these agents influence SB occurrence remains elusive and, based on mouse models, responses to supplements like folic acid vary with the genetic context [34, 84][17, 58, 61]. Although powerful, the mouse is an imperfect surrogate for humans on several counts, among them intergenic regions that differ significantly from the human genome, with less species conservation than protein coding regions. At present, it is not possible to identify maternal-fetal genotypes indicating vulnerability to a teratogenic drug or toxin or to predict which preventive therapy will best ensure healthy pregnancy outcomes for individual couples. There is a pressing need to identify patterns of human SB genetic predisposition, that could lead to better understanding individual prognosis, improved care of SB afflicted infants and enhanced capabilities for birth defect prevention.

Next-generation sequencing offers increasing insight into risk factors for common complex genetic disorders including type II diabetes (1 in 10 in the US[32], schizophrenia (1 in 100) [7, 76] and autism spectrum disorders (1 in 59) [77, 88]. However, less prevalent complex genetic disorders are particularly challenging as they affect relatively small and globally diverse populations (e.g., in the US, 1 in 3000 for NTDs, 1 in 700 for orofacial clefts (OFC) [8] and 1 in 140 for congenital heart disease (CHD) [66]), which requires pooling genetically diverse cohorts that may confound downstream analyses. Genetic studies of the more prevalent structural birth defects like CHD have indicated that, while sequence variants that are common in human populations probably contribute to birth defects, they account for only a small proportion of genetic risk [66], so that genome wide association studies (GWAS) will require thousands of subjects to identify factors. Nevertheless, NTD cases display significant enrichment in rare variants [16, 86], suggesting these changes have greater effect sized and may be identifiable in smaller cohorts, This is the first multicenter SB case-control study to mount a comprehensive, ancestry-matched whole genome sequence (WGS) analysis from a systems biology perspective. This study seeks to identify pathways and biological functions that are disrupted in SB as reflected in their enrichment with genes or regulatory regions harboring rare, likely damaging mutations.

## Results

We obtained WGS data on 310 individuals encompassing 157 SB cases and 153 controls. After quality control screening, remaining samples were analyzed to ensure that no population substructure bias was present. Genetic ancestry was examined and only cases and controls with paired-matched ancestry were selected for further analysis (see Methods for details). Downstream computations were carried out on 298 individuals including 149 cases and 149 non-malformed controls. The mean sequencing depth of the samples was >30X, regardless of their origin (i.e., venipuncture or newborn blood spots) (**Suppl. Table S1, Fig. S1**). Admixture proportions of the selected individuals are shown in **Figure 1A**. Variants were called using a standard pipeline (see Methods).

**Figure 1.**
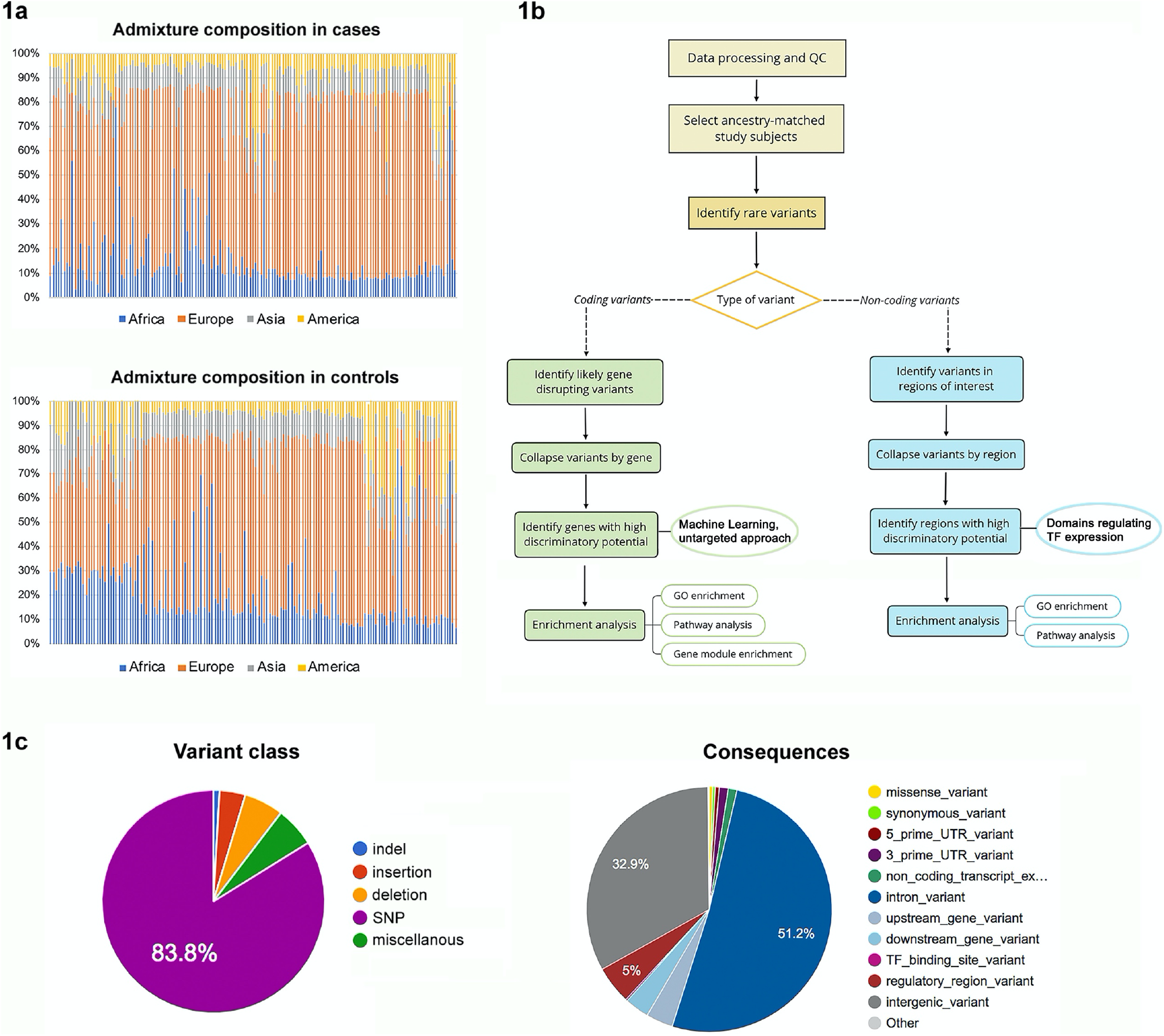
Whole genome sequence analysis overview. **(A)** Admixture composition of the ethnically diverse cohort of 149 spinal bifida cases and 149 ancestry-matched controls used in the analysis. For brevity, the nine gene pools were collapsed by continent. Corresponding gnomAD designations: **(B)** Strategy used to interrogate whole genome sequence data. **(C)** Proportion of variants found in the cohort by type.

Decades of clinical and animal model investigations have led to the consensus that genetic contributions to SB predisposition are most likely due to rare variants, and only a few examples are cited here[15, 41, 52, 87, 89]. However, since some relatively common alleles may contribute to SB genetic risk, we used a somewhat relaxed definition of rare, including for further analysis those variants with an allele frequency (AF) < 0.01 in any population from 1000 Genomes[1], NHLBI GO Exome Sequencing Project (ESP) [80], and the Genome Aggregation Database (gnomAD) [45]. Hence, from a total of 41,005,720 variants at the cohort level, 22,502,019 variants were retained for subsequent steps. **Figure 1B** outlines the workflow used for the genome-wide, ancestry-matched comprehensive set of analyses reported in this manuscript. **Figure 1C** provides a breakdown of the different types of variants found in the study cohort.

### Coding variant analysis supports existing literature and identifies new pathways involving inflammation, innate immunity and cytoskeletal regulation

We previously reported an increased burden of non-recurrent loss-of-function variants in genomes of NTD patients compared to controls, which was consistent across cohorts with different ancestry[16]. Herein, we extended the analysis of coding regions to include all rare likely-gene disrupting (LGD) variants (i.e., frameshift, nonsense, splice donor/acceptor, stop gain/loss and missense predicted deleterious). Thus, 56,210 rare LGD single nucleotide variants (SNVs) and InDels at the cohort level were included in the current analysis. Traditional approaches typically involve finding an association between a variant (or a gene) and the disorder. However, with a limited sample size, this study was underpowered to carry out typical, rare variant association analysis that involves aggregating the effect of rare variants within a gene. More specifically, over 3,000 cases will be required to reach statistical significance at the gene level at 80% power, assuming minor allele frequencies of 5% (see Methods). Accordingly, rare variant aggregate association tests such as SNV-set (Sequence) Kernel Association Test (SKAT-O) [49] did not render any statistically significant results at the single gene level after multiple testing correction.

In view of the complex genetic nature of SB and its relatively low prevalence, systems biology approaches are more appropriate to find statistically significant results after correcting for multiple hypothesis testing. Since the rare (or even private), potentially deleterious variants found in cases are likely to affect different genes within several common pathways or biological processes and functions, we surmised that a machine learning approach can help reduce the genome search space. To test this assumption, the number of rare, LGD protein-coding variants per gene and subject was used as input to determine which of the 13,526 genes harboring these variants were relevant to distinguish between cases and controls (i.e., had high discriminatory potential). Embedded feature selection was employed to perform this process. From the total number of genes used as input to the feature selection step, 6,002 harbored variants in multiple individuals. Random Forests[13] (RFs) were used to build a predictive model of SB, achieving an area under the receiver operating characteristic curve (AUROC) [35]of 0.78 on a completely separate subset of the data (hold-out dataset) never used for training (see Methods). A list of 439 genes was obtained from the full model using embedded feature selection (see Methods, **Suppl. Table S2**). The gene list was then used for enrichment analyses in order to identify pathways, biological processes, molecular functions and cellular components that were overrepresented.

Genes were classified into broad annotation categories as an overview of the biological processes that were affected (shown in **Suppl. Fig. S2**). Interestingly, out of the 439 genes with high discriminatory potential between cases and controls, nine were differentially expressed in a previous transcriptome analysis of fetuses with NTDs[65]. That small study performed genome microarray-based transcription profiling of human fetal amniocyte-derived mRNA from pregnant women at 17-19 weeks’ gestation, comprising seven NTD-affected pregnancies compared to five healthy controls[65] (**Table 1**). Next, we sought to determine whether any of the fetal cortical clusters of genes (“gene modules”) identified in a previously published analysis based on human midgestational (weeks 14-21) RNAseq data[83] were enriched in SB discriminative genes (see Methods). We found that the human fetal gene expression cluster referred to as the “turquoise module” by Walker *et al*. was the only module of six that was significantly enriched in genes with high discriminatory potential in our SB cases (adj. p<0.02) (**Figure 2A**). This turquoise module was described[83] as enriched for specific brain cell types or brain-relevant GO terms involving mitotic progenitors and cell division, and therefore represents an early progenitor class.

**Table 1.** Genes found in this machine learning strategy to have high discriminatory potential between SB cases and controls that were previously found to be differentially expressed in human fetal NTD vs. healthy control amniocytes §.

| Gene | Expression Up/Down | Fold change (log2) | Adjusted p-value |
| --- | --- | --- | --- |
| CGAS* | + | 2.82 | 0.02 |
| GRIN2D** | + | 3.15 | 0.00 |
| MYH11* | + | 2.92 | 0.01 |
| ODF3B | + | 2.69 | 0.04 |
| IVL | - | 2.96 | 0.01 |
| LAMC2* | - | 2.37 | 0.02 |
| SLITRK6** | - | 2.69 | 0.02 |
| USP2* | - | 2.81 | 0.00 |
| ZNF750 | - | 2.23 | 0.04 |

\*=these differentially expressed genes are also found in significantly overrepresented pathways obtained in our analysis
\*\*=these differentially expressed genes have been associated with neuronal synapse assembly and axon pathfinding
§=amniocyte data (log2 fold changes and adjusted p-values) reported by Nagy et al., 2006 (ref. [65])

**Figure 2.**
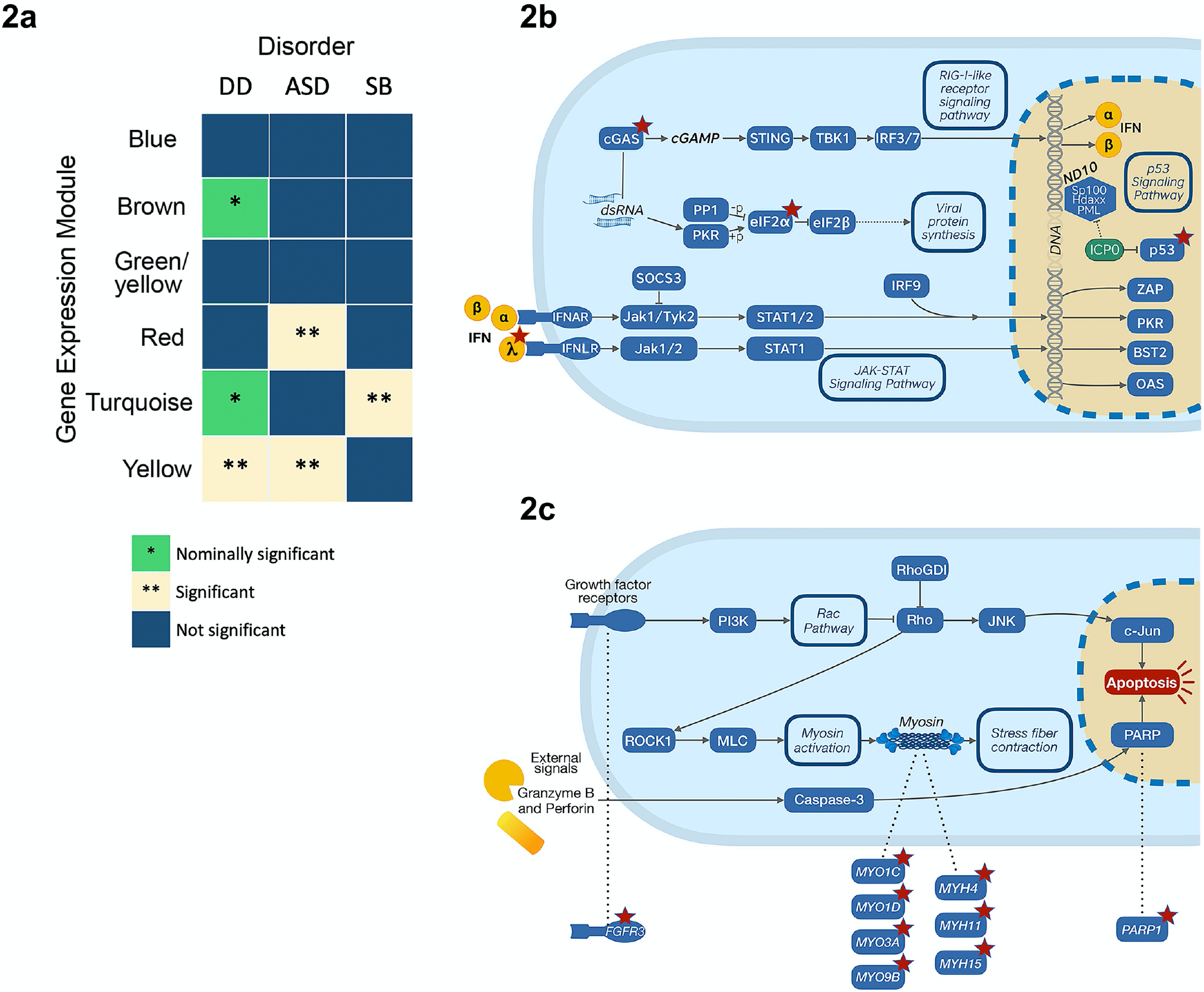
Genes with greatest potential to discriminate between SB cases and non-SB controls and their relationships in pathways. **(A)** Genes with high discriminatory potential to distinguish spina bifida (SB) cases and controls significantly enrich an early progenitor class, gene co-expression module identified in a transcriptome WGCNA study of mid-gestation human cortex[83]. Modules most highly enriched in rare variants found in individuals with developmental delay (DD, neuronal regulation module) or autism spectrum disorder (ASD, neuronal regulation and neurobehavior modules) (Walker *et al*.) are distinct from SB (this study, early neural progenitor proliferation module). **(B)** Pathways related to immunity are enriched with genes that contain likely gene disrupting (LGD) mutations in SB cases and impact the interferon arm of the HSV-1 pathway. **(C)** SB risk genes affect cytoskeletal regulation. Genes enriched with LGD variants in SB cases disrupt RhoGDI signaling and actin-myosin components of the cytoskeleton. Red stars in B and C indicate LGD-enriched genes.

Twenty relevant pathways were overrepresented in genes with the potential to discriminate between SB cases and controls (**Table 2**). Importantly, the pathways with greatest significance were those related to central metabolism (Carbon Metabolism, and Cobalamin Transport & Metabolism, adjusted p-value < 0.001). The variant-enriched genes within these pathways suggest lipid (fatty acid) and glucose metabolism as the aspects most affected in our cohort. This is particularly interesting in that epidemiological data arising after the introduction of folic acid fortification into the US food supply has indicated the persistent risks for NTD may be largely attributable to the rise in obesity and diabetes [40, 56, 62]. This finding provides strong evidence that the proposed approach is pinpointing relevant pathways.

**Table 2.**
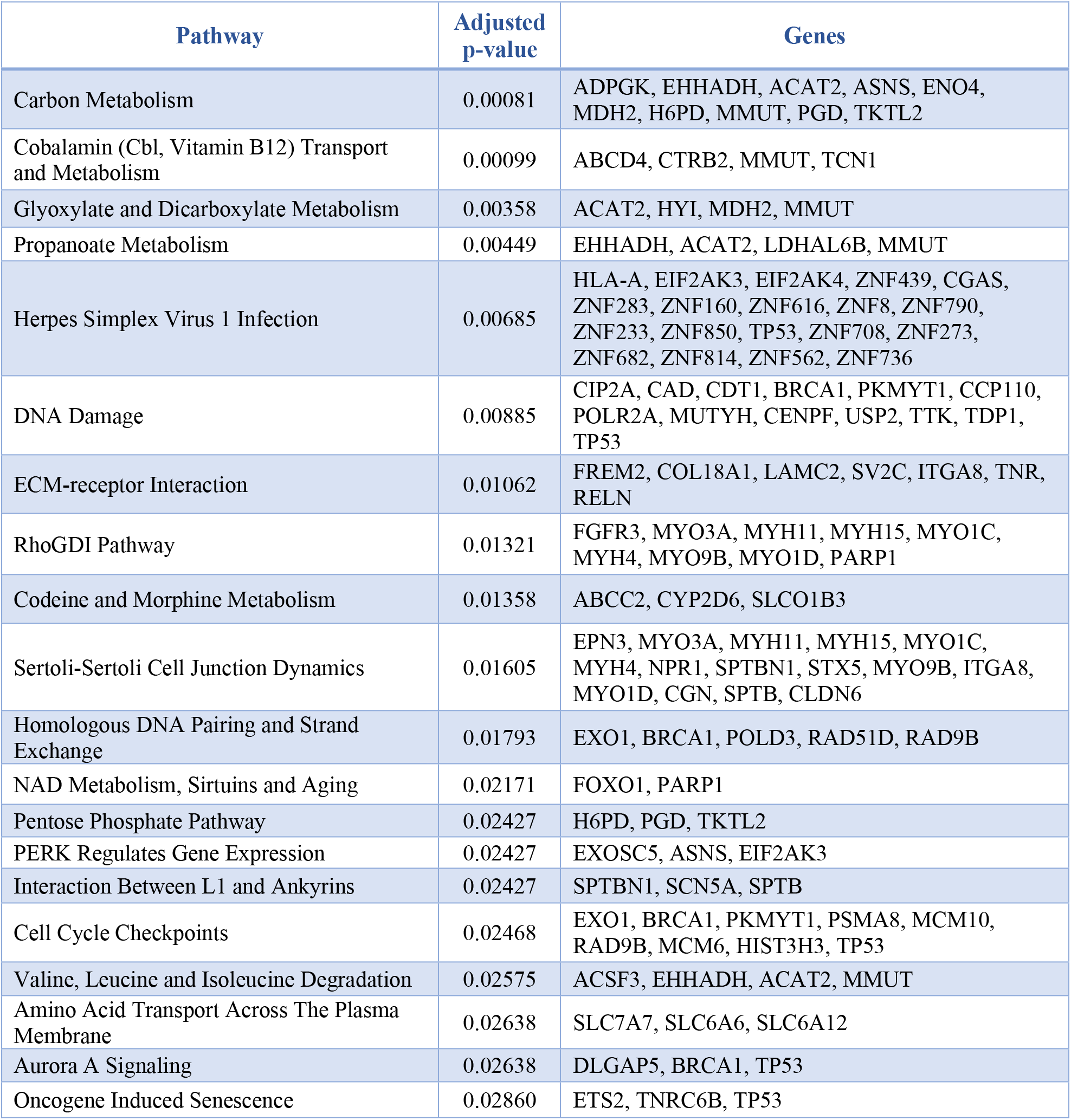
Pathways enriched with genes of high discriminatory potential between SB cases and healthy controls

Additional pathways associated with human SB encompass genes linked to innate immunity and inflammatory response cascades. For example, the Herpes Simplex Virus 1 Infection pathway (**Table 2**, adj. p=0.00685 and **Figure 2B**) includes Cyclic GMP-AMP Synthase (CGAS), whose expression was increased in fetal cells from human NTD cases (**Table 1**). Interestingly, this gene is interferon-inducible and part of innate immunity[14]. The rare LGD variants of the analysis cohort impact genes linked to three critical cascades: RIG-I-like receptor signaling, JAK-STAT signaling and p53 signaling (**Figure 2B**). Another relevant pathway implicated in human SB is associated with response to DNA damage (adj. p=0.00885) and includes USP2, a ubiquitin specific peptidase required for TNF-alpha induced NF-kB signaling. USP2 was also differentially expressed in the fetal cells from NTD cases shown in **Table 1**. Together, these pathways are consistent with previous data implicating immune responses in SB pathogenesis[25] and suggest fetal contributions beyond maternal factors *in utero* produce SB.

The Extracellular Matrix (ECM)-receptor Interaction (**Table 2**, adj. p=0.01062), cytoskeletal regulation (RhoGDI Pathway, **Table 2**, adj. p=0.01321 and **Figure 2C**), and cell-cell signaling (Sertoli-Sertoli Cell Junction Dynamics, **Table 2**, adj. p=0.01605) pathways were also significantly impacted by rare, LGD variant-enriched genes in SB cases. Among these cascades, genes such as Laminin Subunit Gamma 2 (LAMC2) and Myosin Heavy Chain 11 (MYH11) were also found to be differentially expressed in NTD fetal cells (**Table 1**).

Gene Ontology (GO) enrichment analysis showed similar biological processes to be statistically significant (**Suppl. Table S3**). Among these are processes related to cellular/molecular transport (Intraciliary Anterograde Transport, adj. p=0.00008; Amino Acid Transmembrane Transport, adj. p=0.00612), cell migration and morphogenesis (Lateral Motor Column Neuron Migration, Positive Regulation of Trophoblast Cell Migration and Auditory Receptor Cell Morphogenesis, adj. p<0.01), response to stress (EiF2alpha Phosphorylation in Response to Endoplasmic Reticulum Stress, adj. p=0.00081; Negative Regulation of Translational Initiation in Response to Stress, adj. p=0.00315), mitochondrial and nuclear DNA (Mitochondrial DNA Repair and Mitochondrial DNA Metabolic Process, DNA Replication, and Regulation of DNA-dependent DNA Replication Initiation, adj. p<0.005), and one-carbon metabolism (Cobalamin Metabolic Process, adj. p=0.00099; Response to Methotrexate [a folate analog], adj. p=0.00486). Additional results can be found for GO enrichment analysis of cellular components and molecular functions (**Suppl. Table S4 and S5**). LGD variant-enriched genes related to the ciliary base were overrepresented (adj. p=0.03580, Suppl. Table S4), consistent with our previous identification of SB-associated variants in the primary ciliary G Protein-Coupled Receptor 161 (GPR161) that caused mislocalization of the receptor and disrupted downstream signaling[46].

### Non-coding variant analysis points to perturbed core signaling pathways due to the dysregulation of transcription factor genes

When assessing the impact of rare variants in intergenic, non-protein-coding regions it is critical to identify those variants likely to have a deleterious impact on gene regulation, as well as to determine which gene(s) may exhibit altered expression. Enhancers are cis-regulatory elements well-known to modulate gene expression by binding transcription factors (TFs) to facilitate or suppress transcription. Within these enhancer regions, transcription factor binding sites (TFBSs) – short motifs demonstrated to bind TFs – can be affected by single nucleotide changes[71]. Nevertheless, when bound by TFs, enhancers can loop long distances to contact and regulate specific genes; therefore, it cannot be assumed that a rare SNV in a specific enhancer will impact the closest gene. Recent studies elucidate the constraints that restrict each of the ∼1 million documented enhancers in the human genome to specific target gene interactions[3, 42]. Several studies have observed that chromatin loops mediated by CTC-Factor (CTCF) and cohesin bound on both anchors at the loop ends isolate genes from active enhancers, thus leading to a dysregulation of gene transcription units partially or fully within the loop when disrupted[26, 33, 38, 42, 79]

Based on the potential of SNVs to abrogate binding of cognate factors, the non-coding portion of the genome was interrogated, which included 17,548,500 rare non-coding SNVs at the cohort level. Regulatory regions of 106 TF genes previously identified as relatively conserved throughout evolution[6] were analyzed for enrichment in rare non-coding SNVs. Two complementary approaches were used to determine the regulatory region coordinates for these: a more traditional one based on curated enhancer regions (obtained from GeneHancer data available through GeneCards[30]) and a state-of-the-art approach based on CTCF loops (conserved across tissues[55] and mapped during early and later differentiation stages in human embryonic stem cells[42]) (see Methods). Our use of CTCF loop maps to infer the gene being regulated by a rare variant enriched transcription enhancer region is a strategy that leverages critical insights into the 3-D configuration of human ES genomes. This is highly relevant to SB as the neurulation defect arises in or proximal to germinal epithelium within the first 30 to 40 days of gestation, within the typical staging of the maps to which we refer.

The (a) panels in **Suppl. Fig. S3-S6** show the distribution of rare variants in non-coding regions for cases normalized to controls for the coordinates corresponding to each set of coordinates. In this analysis, we identified four TF genes that were enriched for rare SNVs in their regulatory regions in cases compared to controls (**Table 3**). Quantile-quantile (Q-Q) plots, cumulative distribution function (CDF) plots and probability-probability (P-P) plots are also included in supplementary materials (**Suppl. Fig. S3-S6**). **Figure 3A** shows a schematic of the CTCF loops and the distribution within them of those rare non-coding SNVs present only in cases for two TF genes: MAX (myc-associated factor X) and JUND (JunD Proto-Oncogene, AP-1 Transcription Factor Subunit). The CTCF loop associated with MAX contains specifically the cis-regulatory regions in the 3’ end, including the 3’ UTR. Disruption of the 3’ UTR could affect localization, stability, export and translation efficiency of mRNA. On the other hand, JUND’s CTCF loop isolates both the TF and its regulatory regions; hence, disrupting this loop could affect its transcription interactions. In both cases, disruption of regulatory loci within MAX and JUND specific CTCF loops positions the variants to affect expression of these TF genes, in turn to impact the genes those TFs regulate and, ultimately, the pathways in which they are involved.

**Table 3.** Transcription factor genes whose regulatory regions are significantly enriched with rare variants

| **TF** | **Full name** | **Adjusted p-value** | **Coordinates** |
| --- | --- | --- | --- |
| **ZNF274** | Zinc finger protein 274 | 1.64E-11 | GeneHancer |
| **RFX5** | Regulatory factor X5 | 5.25E-05, 7.56E-08 | hESC CTCF loops (naïve and primed) |
| **MAX** | MYC associated factor X | 6.37E-05, 0.018 | CTCF loop conserved across tissues and hESC (naïve) |
| **JUND** | JunD proto-oncogene,<br>AP-1 transcription factor | 0.026 | hESC CTCF loops (primed) |

**Figure 3.**
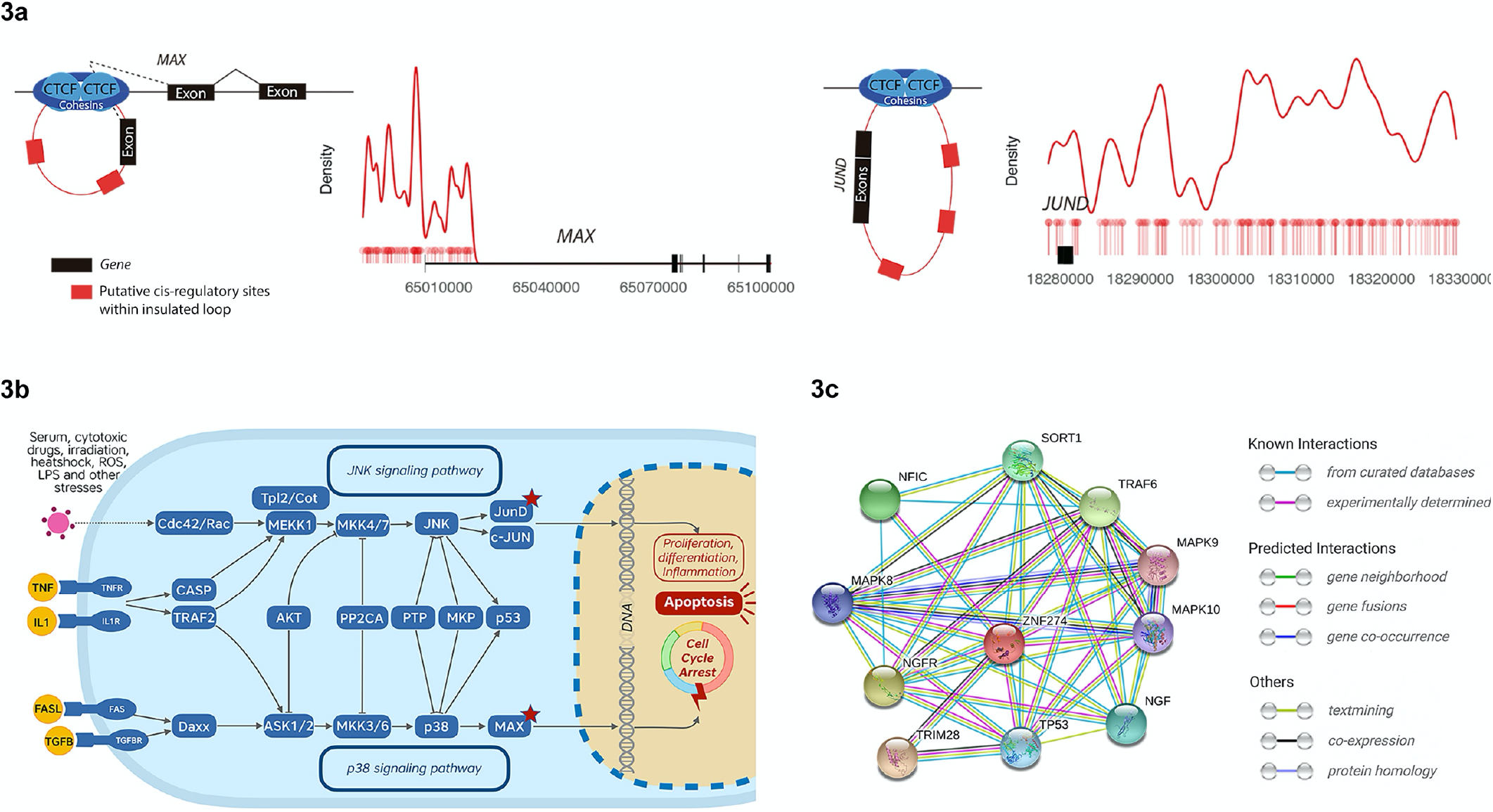
Transcription factor genes whose regulatory regions are enriched with rare non-coding SNVs and their interactions. **(A)** Location of rare non-coding variants within the CTCF loops spanning MAX and JUND in cases. **(B)** Pathways regulating cell processes are impacted by rare non-coding variants. The regulatory regions of MAX and JUND are enriched in rare SNVs, impacting the JNK and p38 signaling pathways. Red stars indicate TF binding site rare variant enrichment in SB cases. **(C)** Interaction partners of ZNF274 based on data from STRING.

Finally, pathway enrichment analysis was carried out using the TF genes identified in the previous analysis (see **Table 3**) as input. Results encompassed several overrepresented signaling pathways pertaining to immunity and the regulation of essential cellular processes, such as cell growth, differentiation and proliferation (**Suppl. Table S6**). As expected, GO enrichment analysis involved biological processes predominantly related to transcription (**Suppl. Table S7**). Nonetheless, it is worth highlighting that terms pertaining to the central nervous system were overrepresented (Response to Axon Injury, adj. p=0.00615; Neuron Apoptotic Process adj. p=0.00807). Additional results can be found for GO enrichment analysis of cellular components and molecular functions (**Suppl. Tables S8 and S9**).

## Discussion

This study comprises a multicenter, population-based, ancestry-matched genome-wide analysis of SB WGS data. Due to the multifactorial nature of SB, our genomic interrogations were stringent, seeking rare changes that produce potentially damaging mutations in protein-encoding sequences or non-coding transcription factor regulatory regions. The strategy pursued here has yielded significant results that stand up to multiple testing correction. Furthermore, the validity of our pathway analysis was supported by two sources of transcriptomic data from human neurodevelopment. First, nine of the genes with discriminatory potential and found in enriched pathways in our study were also differentially expressed in a small survey of differentially expressed mRNAs from mid-gestation fetal amniocytes of NTD affected pregnancies. Second, using our data in a gene module enrichment method (weighted gene co-expression network analysis, WGCNA), we found that variant enriched genes from our SB data overlapped with a gene co-expression module from a study[83] of mid-gestation human cortex, a module that was classified as representing an early progenitor network. This result is appropriate for a structural birth defect that involves germinal epithelium over the first gestational month and supports the relevance of the SB associations detected here.

Among the pathways, defined by PathCards and Gene Ontology, that were most highly enriched with genes that were discriminative for SB were ‘Carbon Metabolism’ and ‘Cobalamin (Cbl, Vit B12) Transport and Metabolism’. It is interesting that the carbon metabolism-related genes found to discriminate SB cases from controls in our cohort are not core players in folate, 1-carbon metabolism, but instead relate to lipid and glucose metabolism. This is particularly relevant in that post-folate fortification epidemiological data has suggested that persistent risks for NTD may be attributable to concomitant population increases in obesity and diabetes [40, 56, 62]. Obesity, metabolic syndrome and diabetes are rising public health concerns both in the US and Qatar populations [5, 62, 74, 81]. This systems biology approach may be particularly suited to detection of physiologically relevant pathways contributing to SB, and may be less subject to ascertainment bias than candidate gene approaches more common in the field.

Another intriguing pathway emerging in this study highlights processes of innate immunity (HSV1 infection and DNA Damage). Among their SB-discriminatory genes, CGAS is known as a sensor of viral dsRNA and DNA damaged dsDNA, it participates in the RIGI-like signaling pathway, and CGAS was differentially expressed in NTD affected human fetal amniocytes [65]. Pathways involving ECM and cytoskeletal regulation mechanisms may illuminate folate resistant mechanisms at work in human NTD, as *Frem2* mutant mice are not protected by folic acid supplementation [58]. The RhoGDI pathway is enriched in several unconventional myosin family members known as regulators of actin-based molecular motors. In particular, MYO1D is necessary for the asymmetric localization of planar cell polarity protein Vangl1[37]. MYO1C serves actin transport to the leading edge of motile cells, while MYO9B is a RhoGTPase activator. Along with the myosin heavy chain MYH gene products, these molecules regulate cell junction dynamics and cytoskeletal contractile elements modulating cell morphologies, and so are positioned to facilitate morphogenetic changes in neural tube cells.

This SB study reaches beyond protein coding sequences to examine nucleotide variation in intergenic functional domains of SB patients. The approach presented here identified four TF genes whose regulatory regions are enriched in variants and are likely contributors to SB risk. Among these, MAX, JUND, and ZNF274 (Zinc Finger Protein 274) stand out (**Table 3**). ZNF274 is a transcriptional repressor involved in epigenetically modified chromatin complexes with SETB1-TRIM28[31] (Figure 3C[78]) and has been associated with p75 neurotrophin mediated signaling (Suppl. Table S6), which participates in key events in spinal cord neuron survival and plasticity[11, 18]. MAX is a bHLH protein a transcriptional repressor acting via the recruitment of a chromatin remodeling complex with histone methylase activity. Among the over-represented pathways encompassing these two TF genes, the MAPK Signaling Pathway (**Suppl. Table S6**) is of particular interest in view of its critical role in brain function[75] and immunity[39]. Both MAX and JUND, through changes in the p38 and JNK Signaling pathways respectively, could disrupt inflammation processes, as well as cell proliferation and differentiation through the cell cycle and induction of apoptosis (**Sup. Table S6, Figure 3B**).

The strength of this approach is in its genome-wide search to identify significant SB associated pathways for hypothesis generation in a manner that avoids cherry picking among genes and pathways already implicated through mouse genetic studies. If limited to SNVs and Indels, our results indicate WGS on some 3,300 SB cases will be needed to establish significance to individual genes. Greater power may be gained from combining gene enrichment by deleterious variants along with damaging structural variation (e.g. CNVs [86]), demanding new computational approaches to accomplish. It is unlikely that a single variant or gene would greatly impact non-syndromic SB risk. However, it is entirely possible that a single pathway could be predisposing if it contained variants affecting multiple genes in the pathway. And only a few LGD containing genes may be necessary to result in SB in an individual, as there are examples in the mouse of NTDs caused by digenic mutations within a pathway. For example, NTDs have been seen in mice heterozygous for mutations in pairs of planar cell polarity pathway genes *Vangl2/Ptk7 [57, 85], Cobl/Vangl2, Vangl2/Scrb*, and *Vangl2/Celsr1* [64], or digenic mutations in cytoskeletal regulators *Enah/Vasp* [60], or cell adhesion genes *Itga1/Itga6* [22]. A case-control study limitation is that it can only illuminate components of SB risk for the affected individual. Determining the recurrence risk for a couple will require WGS investigations of case-parent trios, and these efforts are ongoing. Trio analyses will enable identification of inherited vs. *de novo* variants. Further computational approaches are needed to find potential genetic interactions in individual cases. And genomics will only provide one piece of the complex puzzle that undoubtedly includes epigenetic modifications of genomic DNA and chromatin, often in response to maternal nutrition and/or environmental exposures. As we build population-based genome investigations, it will be important to gather gene expression data from the same subjects—for example from amniocytes of SB affected and control pregnancies.

Hypotheses generated using systems biology-based computational strategies will require functional validation, likely utilizing genome editing in mouse and human stem cell biological systems. These animal and human stem cell models will also offer opportunities to test environmental stressors that mimic toxic exposures and intrauterine conditions that undoubtedly interact with the genome and impact the epigenome to tip the epistatic load toward SB [8]. The approach demonstrated here represents an important step toward an integrated systems view of genetic factors underpinning human neural tube defects. These efforts will undoubtedly have to be combined with investigation of epigenetic, multi-omic, and environmental factors to obtain a full picture required for precision medicine [24]. Importantly, identification of recurrence risk toward new avenues for prevention is only one use of precision medicine. Knowledge of the genetic risk of an individual SB infant—even which pathways are most likely impacted in that individual--could inform prognosis and allow for devising novel early interventions toward optimizing the developmental potential of the child. Systems biology approaches will enable inclusion of relatively rare, complex genetic disorders like spina bifida in this future of 21^st^ century disease prevention and improved individualized healthcare.

## Materials and Methods

### Study subjects

For this case/control study, subjects with non-syndromic SB who displayed myelomeningocele were selected[50, 51]. The initial cohort comprised 310 subjects from different ancestries. Of the 157 SB-affected individuals, 85 were collected in the US and 72 in Qatar. Among the 153 remaining controls are 45 unrelated subjects from the US and 108 unrelated individuals living in Qatar[47].

Human subjects research protocols were approved by Institutional Review Boards (IRBs) in the US (state of California and Stanford University, the University of Texas at Austin, Weill Cornell Medical College-NY), and the Middle Eastern population receiving their healthcare in Qatar (Hamad Medical Corporation and Weill Cornell Medical College-Qatar), including informed consent documentation provided in both English and Arabic.

### Whole genome sequencing

Subject material included genomic DNA extracted specifically for this project from de-identified infant blood spot cards collected from the California Genetic Diseases Screening Program and referred from the California Birth Defects Monitoring Program (CBDMP[20]. Genomic DNA was also derived from venipuncture samples collected from subjects participating in the national Spina Bifida Clinic at Hamad Medical Corporation (Doha, Qatar). Genomic DNA was extracted either from newborn screening bloodspots or infant/child venipuncture samples using the Puregene DNA Extraction Kit (Qiagen, Valencia, California). Input amounts of DNA from infant blood spots were 200-500 ng and inputs from venipuncture samples were 2-3 ug. All DNA samples were whole genome sequenced using Illumina chemistries (v3) on HiSeq2500 instruments to yield short insert paired end reads of 2×100 bp.

### Population structure analyses and case-control matches

Genomic ancestry was calculated from the genotype data by analyzing a set of 130,000 Ancestry Informative Markers reported by Elhaik et al. [28]. Using *supervised ADMIXTURE* [4], we calculated the ancestry of each individual in relation to nine gene pools representing geographic regions around the world (e.g., South Africa) [28]. The output was the admixture proportions of each individual corresponding to those gene pools. To avoid stratification bias due to differences in genetic background, the Pair Matcher (PaM) algorithm was used to genomically match the cases with the controls by their genomic distances [27]. Briefly, *PaM* calculates the genetic distances as the sum of differences between admixture components. The pairing assignments are optimized to maximize the numbers of ancestry matched pairs and ensure that a genetic ancestry-balanced cohort was used in the analysis.

The final study cohort included 298 human subjects. Of the 149 SB-affected individuals, 77 were from the US and 72 from Qatar. Among the 149 ancestry-matched controls were 43 unrelated subjects from the US and 72 unrelated individuals living in Qatar. The remaining 34 controls matching the ancestry of US subjects were selected from the Pan-Cancer Analysis of Whole Genomes (PCAWG) study[19], all of which were germline samples obtained from Caucasian subjects.

### Read mapping, variant calling and annotation

The sequence data were processed using standard pipelines, as described in the Broad Institute’s GATK Best Practices[82]. Reads were aligned to the hg38 reference provided as part of the GATK Bundle using BWA[53]. Variant calling was performed with GATK4[69] and joint genotyping was carried out on the whole cohort, followed by Variant Quality Score Recalibration (VQSR). Quality control (following standard practices such as obtaining sequencing metrics, per sample missing rate and level of heterozygosity), was done to check for DNA contamination and identify outliers, removing those samples with poor quality. Per-variant quality was also assessed and only variants with a “PASS” in the filter column were retained and annotated utilizing Ensembl Variant Effect Predictor (VEP) v.95[59].

### Rare coding variant analysis

Variants in coding regions were filtered to retain only those that are rare (MAX_AF<0.01, as provided by the VEP annotation) in any given population part of 1000 Genomes[1], NHLBI GO Exome Sequencing Project (ESP) [80], and the Genome Aggregation Database (gnomAD) [45]. Next, likely gene-disrupting (LGD) variants were identified as SNVs and Indels, including: 1) loss-of-function variants (LOF; i.e., nonsense, frameshift, splicing, stop gain or stop lost), and 2) missense variants predicted deleterious (by SIFT[67] and/or PolyPhen[2]). Variants meeting the previous criteria (from now on qualifying variants) were collapsed by gene, that is, a matrix with the number of qualifying variants per gene per subject was obtained. A power calculation for individual gene association was obtained using the Genetic Association Study (GAS) Power Calculator[43]. This tool was used to determine the minimum number of subjects required to reach statistical significance at the gene level (p-value=0.0000025). Therefore, assuming a power of 80% and a minor allele frequency (MAF) of 5%, at least 3,300 cases are necessary.

Since the sample size necessary to achieve statistically significant single gene association using rare variant association analysis was well above the available number of cases, an alternative approach was proposed. The matrix of qualifying variants was used as input to a machine learning classifier for embedded feature selection. Hence, genes were selected as part of the learning algorithm, using as class label the group to which each individual belongs (i.e., case or control). A Random Forest (RF) classifier[13] was built for this purpose, using Python’s scikit-learn library[68]. Hyperparameter tuning was performed utilizing *random search* and *grid search* with cross-validation. The final model was further tested on a separate (hold-out) dataset for validation purposes encompassing 20% of the input data. As an additional quality control check, we created RF models on ten sets that were generated by randomly shuffling the group (case/control) labels. This analysis sought to ensure that the model was not learning the noise existing in the data and, as a consequence, would not generalize well.

Features (i.e., the 439 genes with high discriminatory potential) were ranked according to importance, based on the Gini impurity metric. Gini impurity provides a measurement of the likelihood of incorrect classification of a new instance of a random variable, if that new instance was randomly classified according to the distribution of class labels from the dataset) and those with an importance value > 0 were selected for subsequent steps. Genes were next broadly categorized based on GO Slim using WebGestalt[54]. The same genes with high discriminatory potential were used as input to GeneAnalytics[9] for pathway and Gene Ontology (GO) enrichment analyses. Within GeneAnalytics, p-values are calculated assuming an underlying binomial distribution and corrected for multiple comparison using False Discovery Rate (FDR) [10]. Finally, gene module enrichment was carried out as described by Walker *et al*.[83]. Briefly, clusters (gene modules) were obtained by these authors as a result of applying weighted gene co-expression network analysis (WGCNA) [48] to bulk RNA-seq data from mid-gestational (weeks 14-21) human cortex. In the present work, gene module enrichment was calculated employing the same logistic regression model described by Walker and colleagues: is.disease ∼ is.module + gene covariates. P-values were adjusted to correct for multiple testing, applying a Bonferroni correction (as described in the same publication).

### Rare non-coding variant analysis

Variants in non-coding regions were filtered to retain only those single nucleotide variants (SNVs) that are rare (MAX_AF<0.01, as provided by the VEP annotation) in any given population part of 1000 Genomes, ESP, and gnomAD. Regions regulating 106 transcription factor (TF) genes previously identified as relatively relevant during development[6] were obtained. These regulatory regions were defined using data pertaining to curated enhancer GeneHancer data[30] and within CTC-Factor (CTCF) loops spanning each TF gene of interest. Three different sets of coordinates – or catalogs – were used to determine the region coordinates for the CTCF loops, including a dataset of conserved loops across multiple tissues[55], and loops mapped in human embryonic stem cells (hESCs) at an earlier (naïve) and later (primed) developmental stage[42]. CTCF maps from these sources are highly appropriate for this purpose as many CTCF loops are conserved across tissues and developmental stages, and we specifically interrogated those that are known to be conserved. Furthermore, SB arises early in development—before 35 days gestation—and the neural tube is a germinal epithelium, so SB is closely related to hESCs at early (naïve) and more differentiated (primed) progenitor stages.

The subsequent steps were performed for each catalog. First, BEDTools[70] was employed to identify those rare non-coding SNVs that fell within regulatory regions. Similar to the coding variant analysis, variants were summarized following a “regulatory region collapsing” strategy. To identify regions with high discriminatory potential, regulatory regions associated to a TF gene were tested for enrichment in cases vs. controls. For this purpose, the proportion of SNVs in cases divided by controls was calculated and the *fitdist* function within the fitdistrplus R package[23] was used to determine which regulatory regions were significantly enriched. P-values were FDR adjusted to correct for multiple comparisons.

Finally, the list of TF genes whose regulatory regions were significantly enriched with SNVs (FDR<0.05) in at least one of the catalogs was used as input to pathway and GO enrichment analysis. Similar to the coding variant analysis, this was carried out employing GeneAnalytics.

## Supporting information

Supplementary Data A-Pulido et al.

## Acknowledgments

We thank Ms. Amira Assad, Project Coordinator at WCM-Q, for invaluable efforts toward patient enrollment. We thank the California Department of Public Health Maternal Child and Adolescent Health Division for providing data. The findings and conclusions herein are those of the authors and do not necessarily represent the official position of the California Dept. of Public Health.

## Notes

### Competing Interest Statement

Dr. Finnell formerly held a leadership position with the now dissolved TeratOmic Consulting LLC. He also receives travel funds to attend editorial board meetings of the Journal of Reproductive and Developmental Medicine published out of the Red Hospital of Fudan University. Dr. Elhaik consults for the DNA Diagnostics Center (DDC). The remaining authors have no competing interests.

