## Supplementary Data A-Pulido et al. for "Systems Biology Analysis of Human Genomes Points to Key Pathways Conferring Spina Bifida Risk"

**This PDF file includes:**

Figures S1 to S6

**Other supplementary materials for this manuscript include the following:**

Data Excel Tables S1 to S9

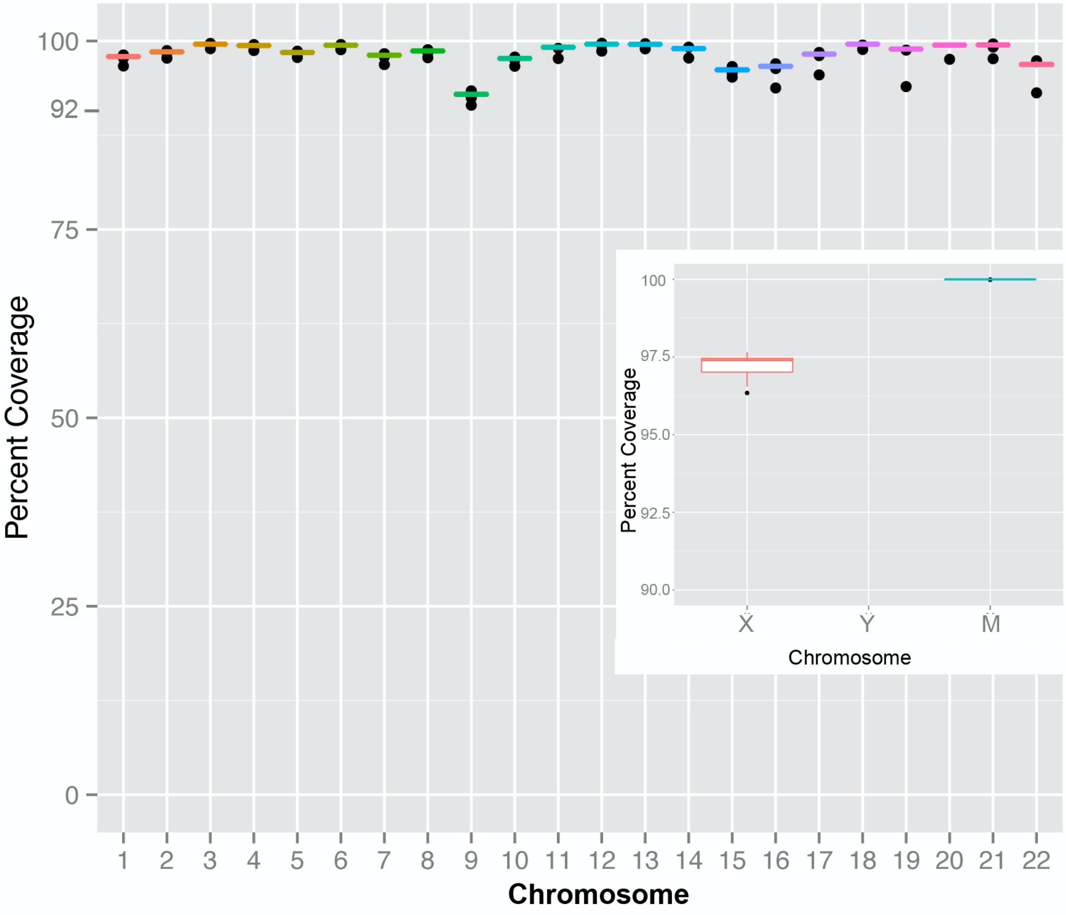


**Figure S1a.** Survey of genome coverage by chromosome for the WGS cohort sequenced in this study. M=mitochondrial DNA

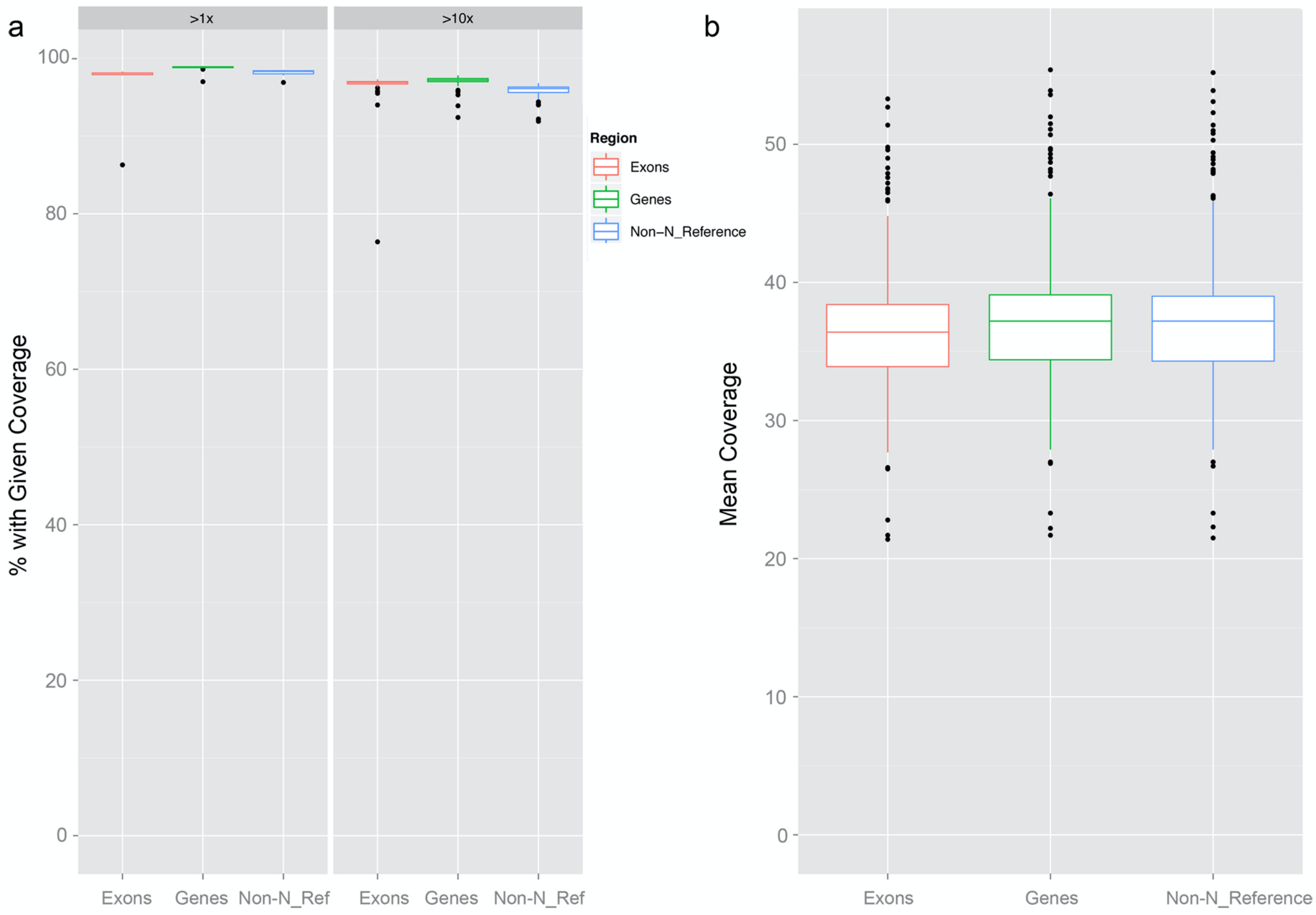


**Figure S1b**. Survey of depth of coverage over exons, genes and whole genome (Non-N_reference sequences) for the WGS cohort sequenced in this study. **(a)** Percentage of samples with more than 1X or 10X coverage. **(b)** Mean coverage depth (37-38X, with none less than 20X).

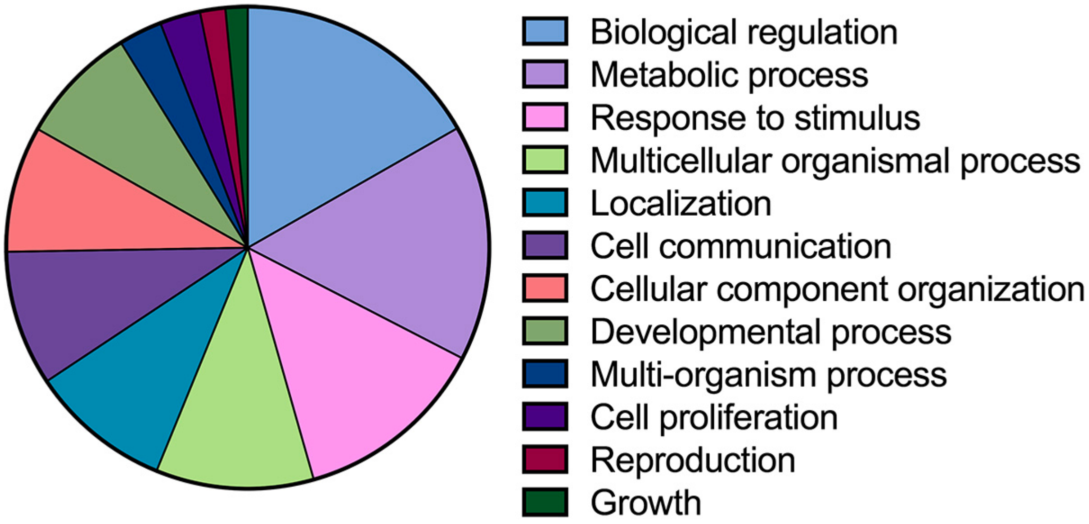


**Figure S2.** Categorization (GO terms) of genes with high discriminatory potential between cases and controls. Genes that could potentially confer risk of developing SB are classified into broad categories based on biological function.

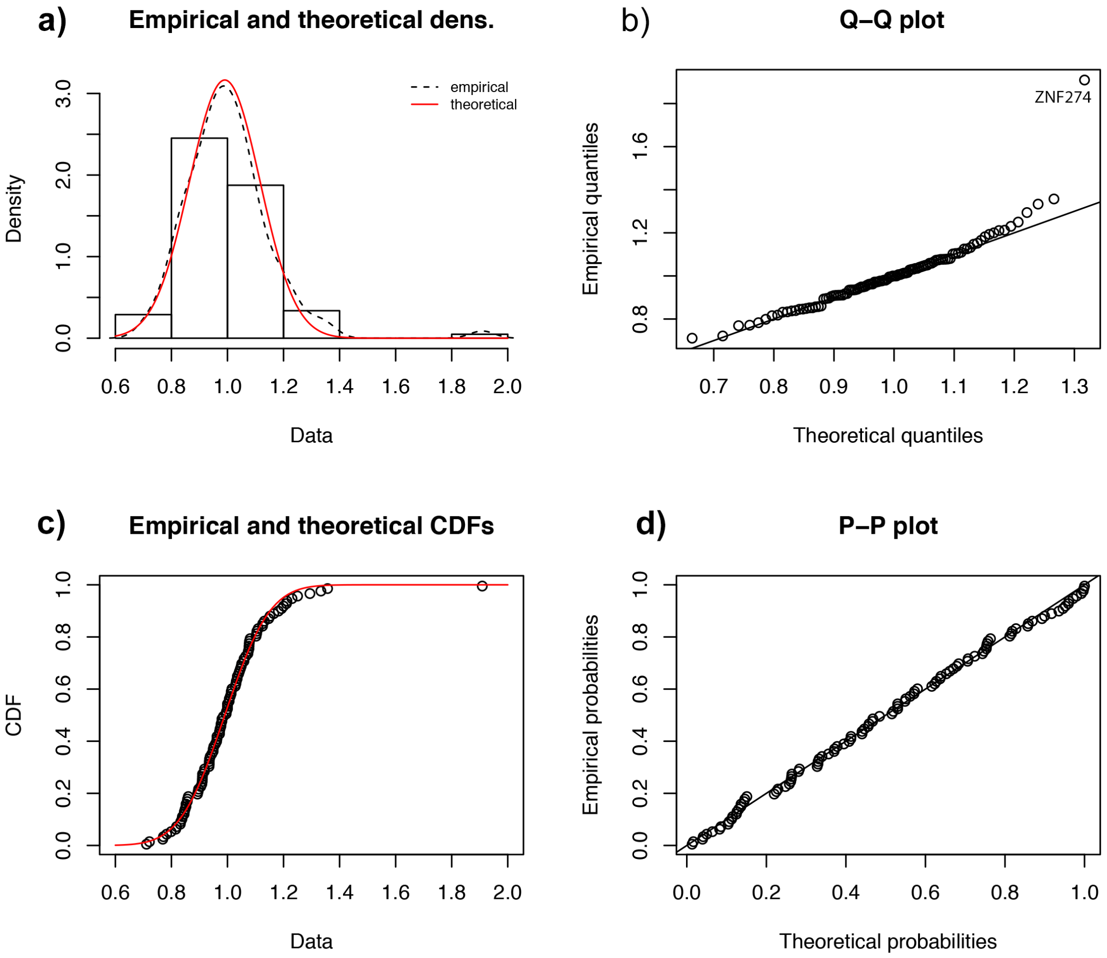


**Figure S3**. Results of the are non-coding analysis using **GeneHancer coordinates**, including: **(a)** distribution of variants, **(b)** quantile-quantile plot, **(c)** cumulative distribution of variants and **(d)** probability-probability plot.

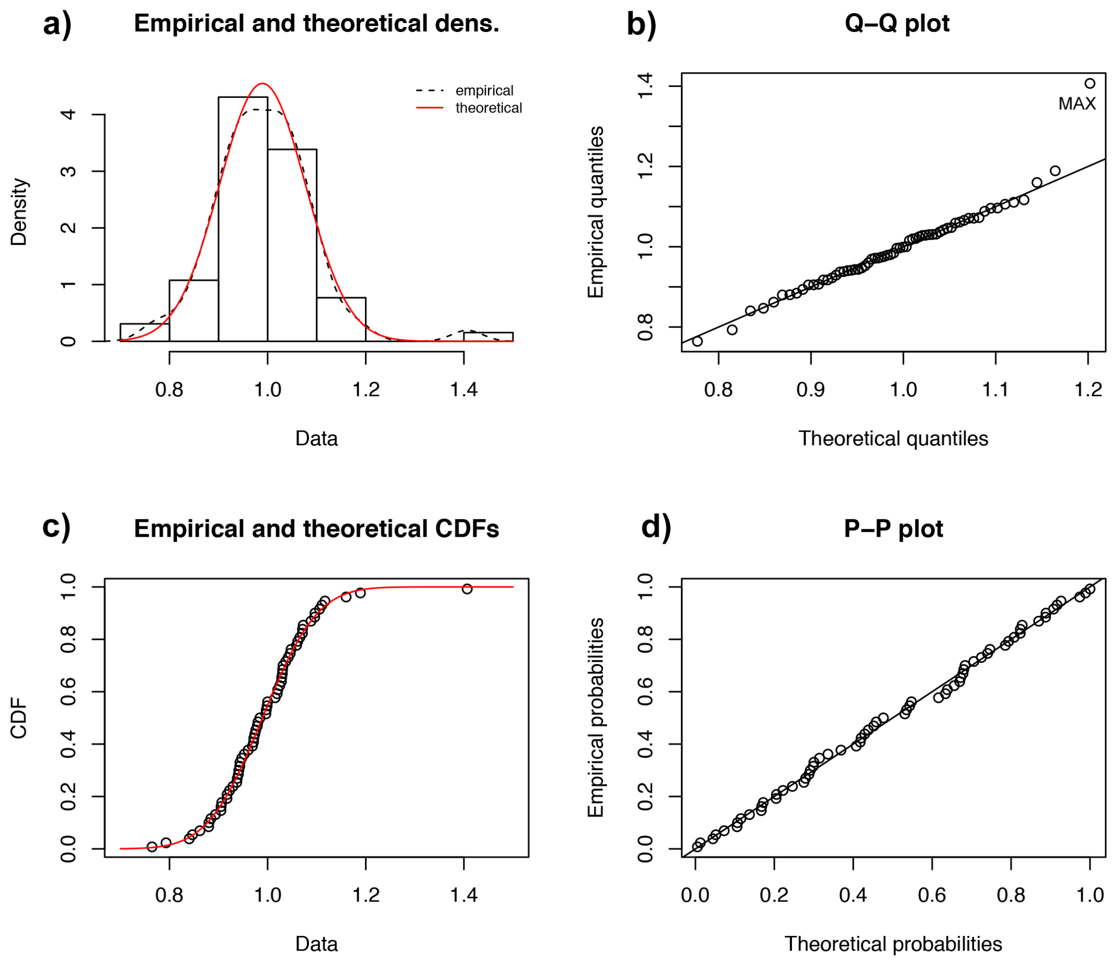


**Figure S4**. Results of the are non-coding analysis using **coordinates for CTCF loops** conserved **across tissues**, including: **(a)** distribution of variants, **(b)** quantile-quantile plot, **(c)** cumulative distribution of variants and **(d)** probability-probability plot.

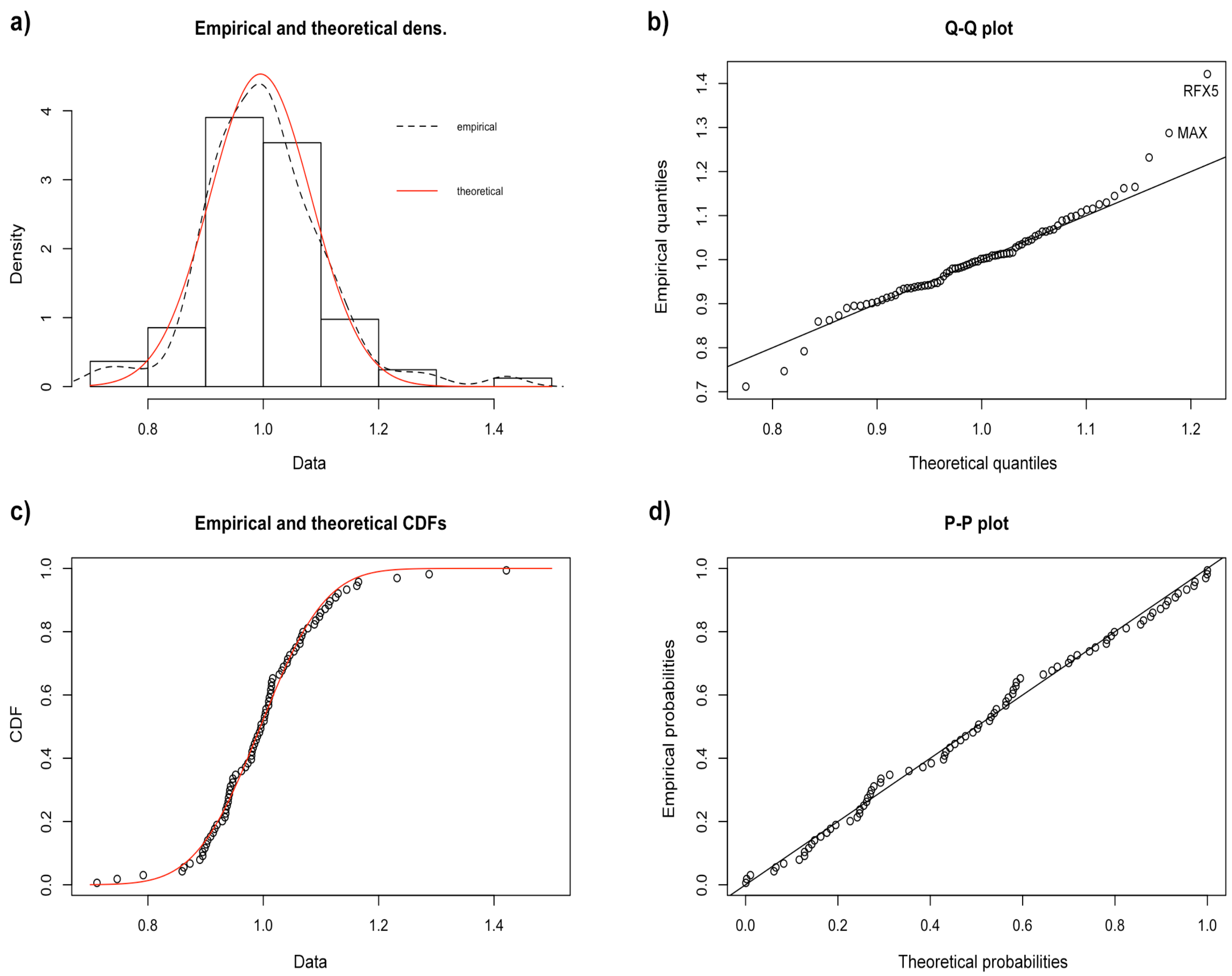


**Figure S5**. Results of the are non-coding analysis using **coordinates for CTCF loops** conserved in **naïve hESCs**, including: **(a)** distribution of variants, **(b)** quantile-quantile plot, **(c)** cumulative distribution of variants and **(d)** probability-probability plot.

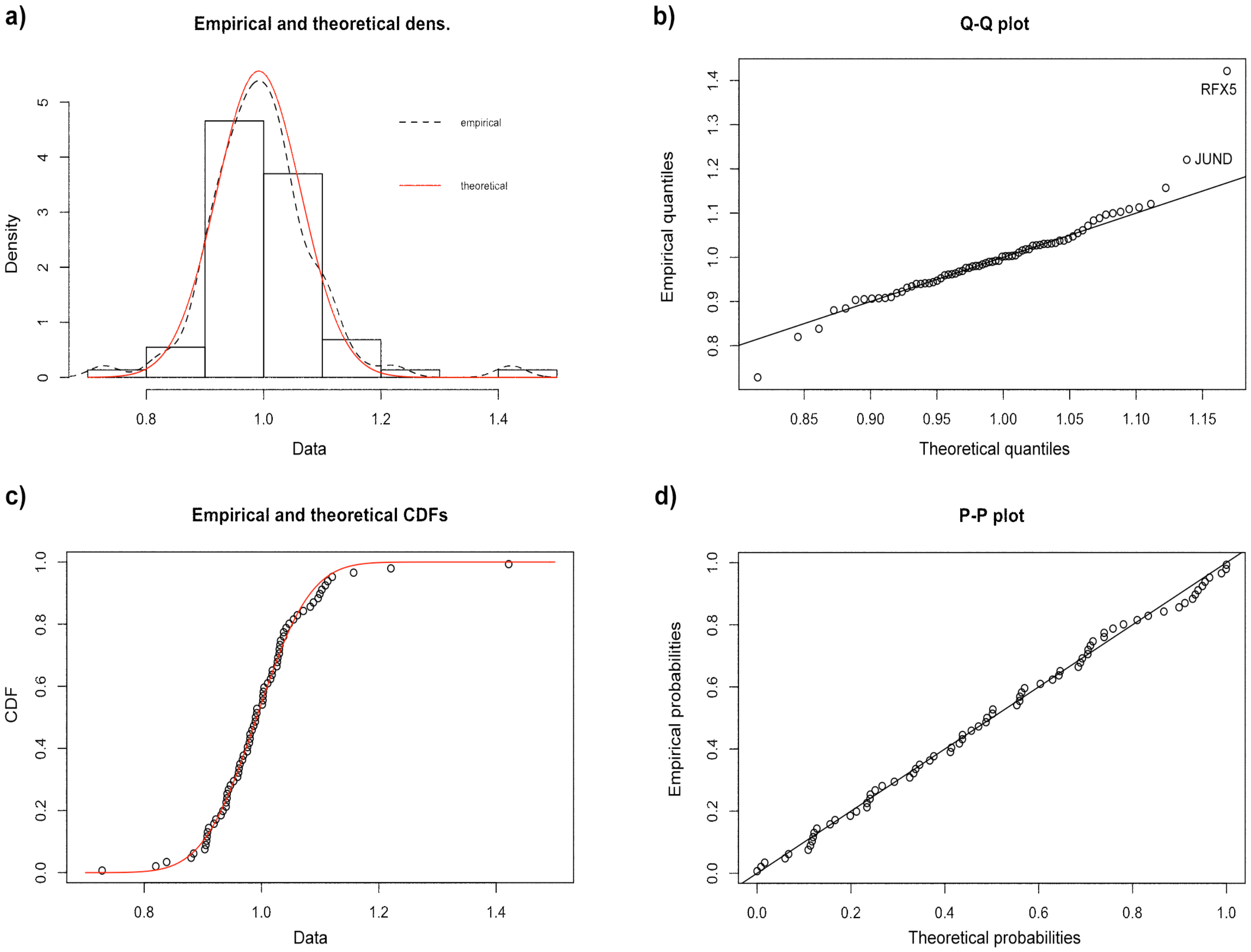


**Figure S6**. Results of the are non-coding analysis using **coordinates for CTCF loops** conserved in **primed hESCs**, including: **(a)** distribution of variants, **(b)** quantile-quantile plot, **(c)** cumulative distribution of variants and **(d)** probability-probability plot.

**Data S1. (separate file)**
Summary of coverage depth for WGS cohort sequenced in this study.

**Data S2. (separate file)**
Genes with rare, LGD variants that show high discriminatory potential between cases and controls.

**Data S3. (separate file)**
Biological processes enriched with rare LGD variants (based on GO enrichment analysis).

**Data S4. (separate file)**
Cellular components enriched with rare LGD variants (based on GO enrichment analysis).

**Data S5. (separate file)**
Molecular functions of genes enriched with rare LGD variants (based on GO enrichment analysis).

**Data S6. (separate file)**
Pathways enriched with TF genes whose regulatory regions encompass significantly more rare variants in SB cases than controls.

**Data S7. (separate file)**
**Biological processes** served by **TF genes** whose Transcription Factor Binding Sites are enriched with rare non-coding SNPs (based on GO enrichment analysis).

**Data S8. (separate file)**
**Cellular components** affected by **TF genes** whose regulatory regions are enriched with rare non-coding SNPs (based on GO enrichment analysis).

**Data S9. (separate file)**
**Molecular functions** served by **TF genes** whose regulatory regions are enriched with rare non-coding SNPs (based on GO enrichment analysis).
